# MOLECULAR INSIGHTS INTO BINDING BEHAVIOUR OF LAMOTRIGINE WITH INITIATION FACTOR 2 PROTEIN: AN INTEGRATED COMPUTATIONAL STUDIES

**DOI:** 10.1101/2022.08.26.505506

**Authors:** Smriti Arora, Jeevan Patra

## Abstract

**Aims:** The ribosomal protein (r-protein) of bacteria is composed of 2.6 MDa ribonucleoproteins of the 30S and 50S subunits, which are essential elements for protein translation. The translational initiation step is an intensive regulated multi-step reaction in protein biosynthesis. During bacterial protein synthesis, the correct reading frame of the mRNA defines when the initiator fMet-tRNAiMet binds to the start codon AUG at the P-site of the 30S subunit. The formation of the P-site of the 30S subunit initiation complex (30S-IC) is governed by three ubiquitous initiation factors (IFs) such as IF1, IF2, and IF3. IF2 protein is an essential player that plays during the last stage of the initiation process. Earlier, Stokes and his co-workers studied chemicals probes using 30K diverse drugs that induced cold-sensitive growth inhibition in the bacterium. The assay studies revealed, Lamotrigine (LTG) effectively binds at domain II of IF2 protein. In our research, we took an attempt in identifying promising active residues that could responsible for anti-bacterial bioactivity with help of computational studies.

**Computational Methods:** In the present study, initially, we performed C-α backbone alignment with the retrieved IF2 chain from AlphaFold. Further, we utilized SiteMap and CastP for the identification of plausible active binding sites. Further, we bound LTG with the designated domain(s) of IF2 protein and studied its binding affinity potential with help of adaptive molecular dynamics simulations at atomic levels using Desmond.

**Key Findings:** Our research findings have shown accurate results and we could able to prove the assertion in contrast with the findings of Stokes and his co-workers where the LTG bind at domain II of IF2 protein. The key interacting residue Glu179 was revealed to have strong hydrogen bonding contacts with LTG at the sub-nanomolar range. In addition, we predicted the alternative promising site I Further, we gained in-depth analysis for studying multiple sites, to understand the synergism inhibitory activity. Promisingly, LTG could be able to bound with at Site 1 showing better affinity over the proposed domain II and other predicted sites. The adaptive molecular dynamics studies confirmed the promising active residues

**Significance:** The binding site predictions approach provides an insight for further development of anti-bacterial therapeutics that might helpful for bacteria disease management and exhibiting inhibitory activity against various strains.

## BACKGROUND

The ribosomal protein (r-protein) of bacteria composed of 2.6 MDa ribonucleoproteins of 30S (small) and 50S (large) subunits, which are essential elements for protein translation. The structural and functional understanding of these ribosomal translation remains to be mysterious (Moore, 2012). During the process of *Escherichia coli* growth, cellular energy consumptions performed by ribosome biogenesis, along with other processes relied within it such as translation, modification, folding and binding of r-protein and releasing of ribosome biogenesis factors (Maguire Bruce, 2009). Inside the molecular cellular level, ribosome biogenesis factors (approx. sixty factors) endured in binding with ribosomal particles to enhance its efficiency of matured subunits (Bunner et al., 2010). Ribosome biogenesis factors are the club of proteins that are assembled in 3D jigsaw that supports to assemble fifty proteins and three RNAs. Moreover, these biogenesis factors prevent the entry of immature subunits into the translational processes (Lebaron et al., 2012; Strunk et al., 2011; Strunk et al., 2012). Several conventional strategies such as genetic perturbation, and genetic in-activation was implemented to probe the proteins functions. However, few drawbacks and difficulties of creating conditional alleles are the pitfalls with such conventional strategies.

Translational initiation step is an intensive regulated multi-step reaction in protein biosynthesis (Marintchev & Wagner, 2004). Generally, the protein biosynthesis occurs in four phases – initiation, elongation, termination and recycling controlled and regulated by ribosome and translational guanosine triphosphatase (trGTPase) respectively. During bacterial protein synthesis, the correct reading frame of the mRNA defines when the initiator fMet-tRNA_i_^Met^ binds to the start codon AUG at the P-site of 30S subunit. The formation of P-site of 30S subunit initiation complex (30S-IC) governed by three ubiquitous initiation factors (Ifs) such as IF1, IF2 and IF3. From these three IFs, translational guanosine triphosphatase (GTPase) of IF2 is an essential player plays during the last stage of the initiation reaction (Simonetti et al., 2009). The IF2, associates 30S-IC with the 50S ribosomal subunit for the formation of 70S-IC. The structural insights of IF2 comprises five domains such as conserved N-terminal domain and C-terminal domains I-IV (Gualerzi et al., 2001). The crucial features of C-terminal domain I is to confer GTPase activity, domain II and III facilitates to bind with the 30S subunit (Simonetti et al., 2013) and domain IV promotes the contacts with the fMet-tRNA_i_^Met^ initiator (Laursen et al., 2005; Spurio et al., 2000). Earlier deeper understanding on the structural insights were gained from the X-ray crystallography IF2 structure excluding domain IV (Eiler et al., 2013; Simonetti et al., 2013) and low resolution cryo EM that elucidated overall mapping of IF2 between the ribosomal subunits (Allen et al., 2005; Julián et al., 2011; Marshall et al., 2009; Simonetti et al., 2008). Sprink and his coworkers, determined the cryo-EM IF2 structure (3.7Å resolution) complexed with non-hydrolyzable guanosine triphosphate derivative and fMet-tRNA_i_^Met^ initiator in the *Escherichia coli* bacteria (Thiemo Sprink et al., 2016). In addition, mechanistic understanding at the final steps of translation initiation studied with the intrinsic confirmational modes of 70S interplaying in between the IF2 and transfer RNA (tRNA) with the ribosome.

Small molecules (less than 300 MW) probing against target candidates widely gained interest across scientific communities and emphasized in the field of drug discovery paradigm. These molecules can be easily studied at the drug target inhibition thus considered to be an elegant probe for protein functions. In addition, inhibiting ribosome biogenesis with help of probing small inhibitors could justify as the promising tool for controlling complex process such as protein assembly factors. Studies shown the antibiotics could be the best probing efforts to understand protein translations. So far very limited research been studied on identification of small chemical probes that could inhibit bacterial ribosome in protein translation. IF2 protein involved in the production of other proteins which might having the role in ribosomal assembly. To best of our knowledge and intensive literature exploration, Stokes and his co-workers studied chemicals probes using 30K diverse drugs those induced cold sensitive growth inhibition in the bacterium (Stokes et al., 2014). From their research study, cold sensitive lamotrigine (LTG) an anticonvulsant drug could able to abort the ribosomal factors assembly. LTG, accumulates the immature 30S and 50S ribosomal subunits at cold temperature (15°C). The assay studies and the structural feature at the domain II revealed that the LTG promisingly shown inhibition towards IF2. However, spontaneous mutation suppressor blocks and makes LTG incapable in perturbing protein synthesis. In this study, we identified putative active residues at domain II and performed best site predictions for the LTG binding surface pocket based on the reported cryo-EM structure. Further, we performed adaptive dynamics studies to proven the assertion governed by the Stokes and his team.

## COMPUTATIONAL METHODS

### Protein structural alignment and preparation

To identify promising residues where LTG could bind for its inhibition activity, we retrieved cryo-EM structure PDB ID 3JCN (Thiemo Sprink et al., 2016), with having resolution of 4.60Å, from RCSB Protein Data Bank (https://www.rcsb.org/structure/3jcn). Protein structure was prepared using protein preparation wizard in Maestro panel. To prepare the structure, the protein preparation module was used to add the hydrogen atoms, remove the waters beyond 5 Å from the binding sites, and optimize the structure for creating an H-bond network. Finally, the energy was minimized using optimized potentials for liquid simulations (OPLS3e) force field with a default setting of 0.30 Å root mean square deviation (RMSD) using imperf script. The overall stability and stereochemical quality of the protein in the 3D structure were determined using the Ramachandran plot.

### Prediction of binding sites and grid generation

Before performing ligand docking, the identification and characterization of the active binding site should be carried out for a more reliable and accurate molecular docking. The active sites for target-ligand binding interaction were predicted using the SiteMap module of Maestro molecular modeling platform v12.8 by Schrödinger program. SiteMap can determine the active binding sites in a large-scale validation with best results for the sites that can bind ligands with the sub-nanomolar association, and in addition, it can propose an adjusted version of the score that accurately classifies the druggability of proteins (Halgren, 2009). The sites predictions of 4 Å were considered with maximum of best 10 iterative sites. The shallow binding pockets were enabled to help us to analyse the potential of druggability score according to the volume and size score of the predicted pockets. All sites were selected with 10 site-pointing groups and 15 individual reported sites. The hydrophobicity of the predicted sites selected to the less reactive with the use of standard grid. All the resulting five binding sites were studied thoroughly for further molecular docking (equation). The parameters are size, volume, degree of enclosure/exposure, degree of contact, hydrophobic, hydroiphilic and hydrogen-bonding (both acceptors and donors).

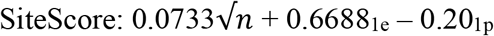

Where, n: the number of site points (capped at 100), e: enclosure, p: hydrophilicity of the site (capped at 1.0).

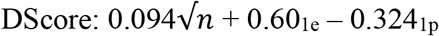

The SiteMap suggest that a cut-off of 0.80 that used to differentiate between drug-binding and non-drug-binding sites, with scores more than 1.0 being indicative of highly promising sites. The Dscore can help to distinguish between undruggable and druggable sites (Halgren, 2009; Wang et al., 2005).

The receptor grid was developed using Glide (Glide, Schrödinger, generated LLC, New York, NY, 2020) by preferring the predicted active sites. The box was centroid of the active site of the protein receptor with the default parameters of van der Waals scaling factor of 1 Å, partial charge cut-off at 0.25, and docked ligand length of 10 Å.

### Ligand Preparations, Molecular Docking and Visualizations

The wet-lab studied drug lamotrigine were retrieved form the PubChem in sdf file format and subjected to LigPrep for the ligand preparations. The LigPrep were designated with using the forcefield of OPLS3e and the generated ionization using Epik and 7.0 ± 2.0 pH. The desalt and tautomers were enabled for performing and yielding the computation that can retain specified chiralities utmost 32 per ligand. Molecular docking is a structure-based drug design approach to identify the essential amino acid interactions between the selected protein and generated ligands with low energy conformation. Minimum interaction of the ligands characterized by the scoring function which used to foretell the binding affinity with the receptor. Glide Standard precision (SP), docking protocol was applied without smearing any constrain. Flexible docking with Glide Standard precision (SP) protocol was performed to predict the binding affinity and ligand efficiency as inhibitor of IF2 drug target. The docking simulation studies were assigned using the standard precision (SP) keeping ligand flexibility of ring and nitrogen orientations and the bias sampling of all torsions at predefined functional groups. Concluding energy assessments were performed with the dock score. All visualizations studies were performed with help of UCSF Chimera and PyMol (DeLano, 2002; Pettersen et al., 2004).

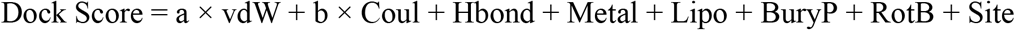

where, a and b are co-efficient constant for vdW and Coul, respectively. vdW = van der Waals energy; Coul = Coulomb energy; Hbond = Hydrogen bonding with receptor; Metal = Binding with metal; Lipo = Constant term for lipophilic; BuryP = Buried polar group penalty; RotB = Rotatable bond penalty; Site = active site polar interaction.

### Molecular Dynamics Simulations

The molecular dynamics simulations were performed up to 100ns using Desmond package (Bowers et al., 2006; Chow et al., 2008). Both the wild and mutant complexes were placed in orthorhombic box at absolute size and solvated using Transferrable Intermolecular Potential with 3-points (TIP3P) using Desmond system builder (Mark & Nilsson, 2001). The simulation system was neutralized with counter ions and salt concentration of 0.15M NaCl. All complex were described with the OPLS force field that were assigned to each simulation run for 100ns. The dynamics simulation was maintained with the isotropic Martyna-Tobias-Klein barostat and Noose-Hoover thermostat at 1 atm and 300K pressure and temperature respectively (Martyna et al., 1992; Martyna et al., 1994). The smooth Particle Mesh Ewald method used for evaluating long range coulombic interactions with a short-range cut-off at 9.0 Å (Essmann et al., 1995). The binding free energies of all protein-protein complexes with individual residue free energy contributions were computed and estimated with help of Molecular Mechanics Generalised Born Surface Area (MM-GBSA) using HawkDock (Weng et al., 2019). All the pictorial representation was drawn and retrieved using Visual Molecular Dynamics – 1.9.3 (Humphrey et al., 1996).

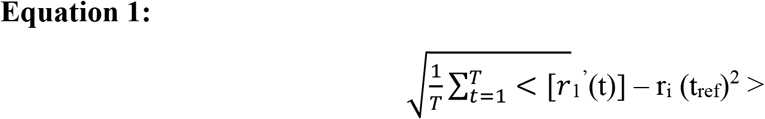

Where, T is the trajectory time over which the RMSF is calculated, t ref is the reference time, r i is the position of residue i; r’ is the position of atoms in residue i after superposition on the reference, and the angle brackets indicate that the average of the square distance is taken over the selection of atoms in the residue.

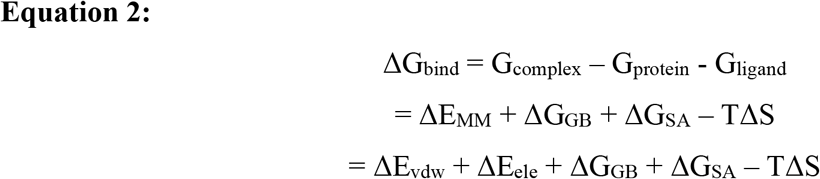

Where, ΔE_MM_ is the interaction energy with two phases ΔE_vdw_ (van der waals energy) and ΔE_ele_ (electrostatic energy) (Genheden & Ryde, 2015). ΔG_GB_ and ΔG_SA_ are the de-solvation free energy (Onufriev et al., 2004). Normal mode analysis used for evaluating entropy contribution to the binding free energy.

## RESULTS AND DISCUSSIONS

### Protein Sequence Alignment and Protein Preparation

Initially, we retrieved the IF2 protein PDB 3JCN of *E. coli* organism from the UniProt database. From the structural insights, it revealed that the IF2 protein having missing residues (1-381). With the support of AI predicted Protein database AlphaFold (Jumper et al., 2021; Varadi et al., 2021) we retrieved IF2 protein with having whole sequence of 890 residues - POA705 (figure 1A) (https://alphafold.ebi.ac.uk/entry/P0A705) for C-α backbone. The powerful AI predicted AlphaFold have predicted the missing residues and cross validated with help of the predicted alignment error shown (Figure 1B).

**Figure 1:**
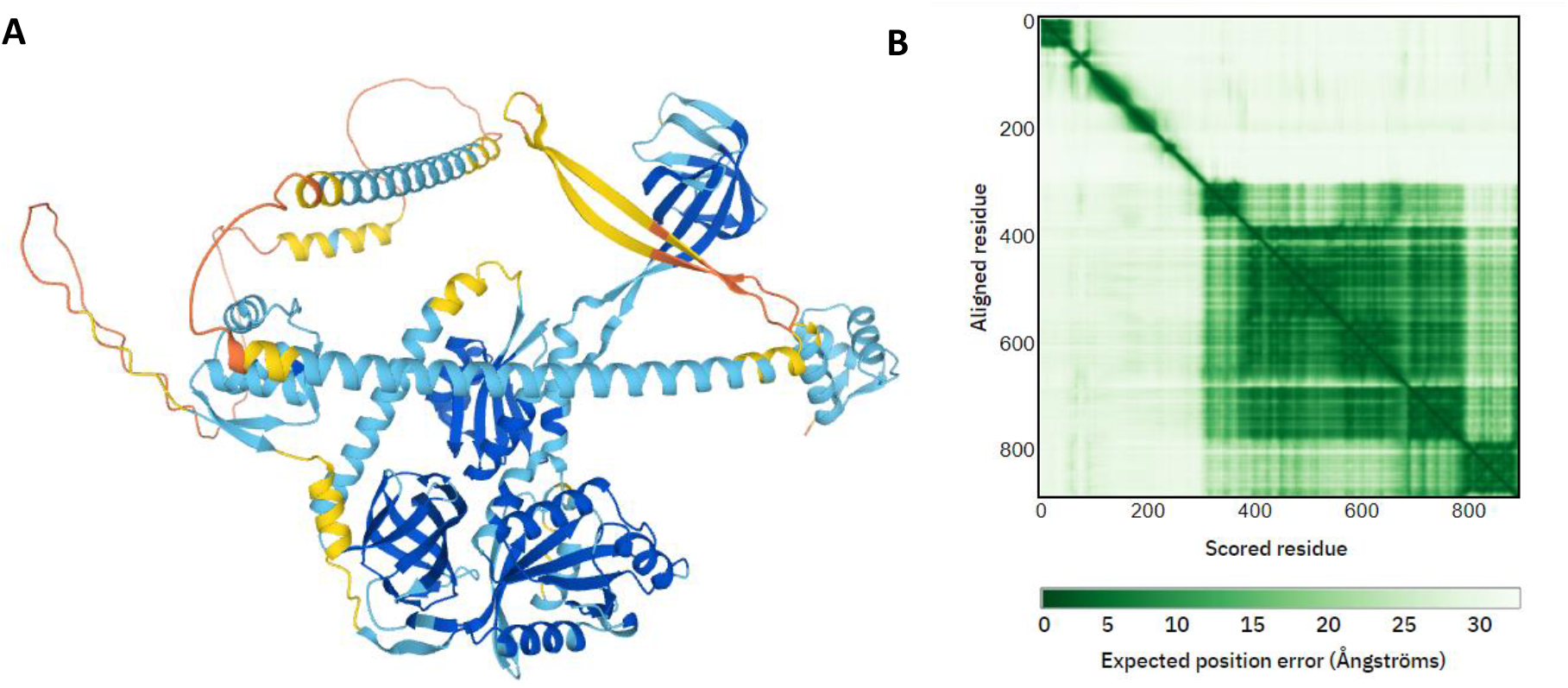
Molecular representation of (A) IF2 protein (POA705) from the AlphaFold database with (B) predicted aligned error

The retrieved protein POA705 from AlphaFold having full 890 residues were structurally aligned. The C-α backbone of proteins were aligned with the Atomic Sequence Alignment (ASL). The resultant protein sequence having RMSD of 2.072 Å and align score of 0.171 (Figure 2). The newly constructed IF2 protein was coordinated precisely and subjected for full protein preparations. The polar receptors assigned bond orders using CCD database, zero order to bond metals and disulphide bonds and generated hetero-states using Epik. The H-bond assignment were performed using sample water orientations using PROPKA at neutral 7.0 pH. Further the assigned H-bond were optimized and restrained minimization to 0.30 Å using OPLS3e force field. The overall protein stability and stereochemical quality in the 3D structure were determined using the Ramachandran plot. This confirms us the predicted alignment error of AlphaFold of IF2 protein was accurate for further informatics studies.

**Figure 2:**
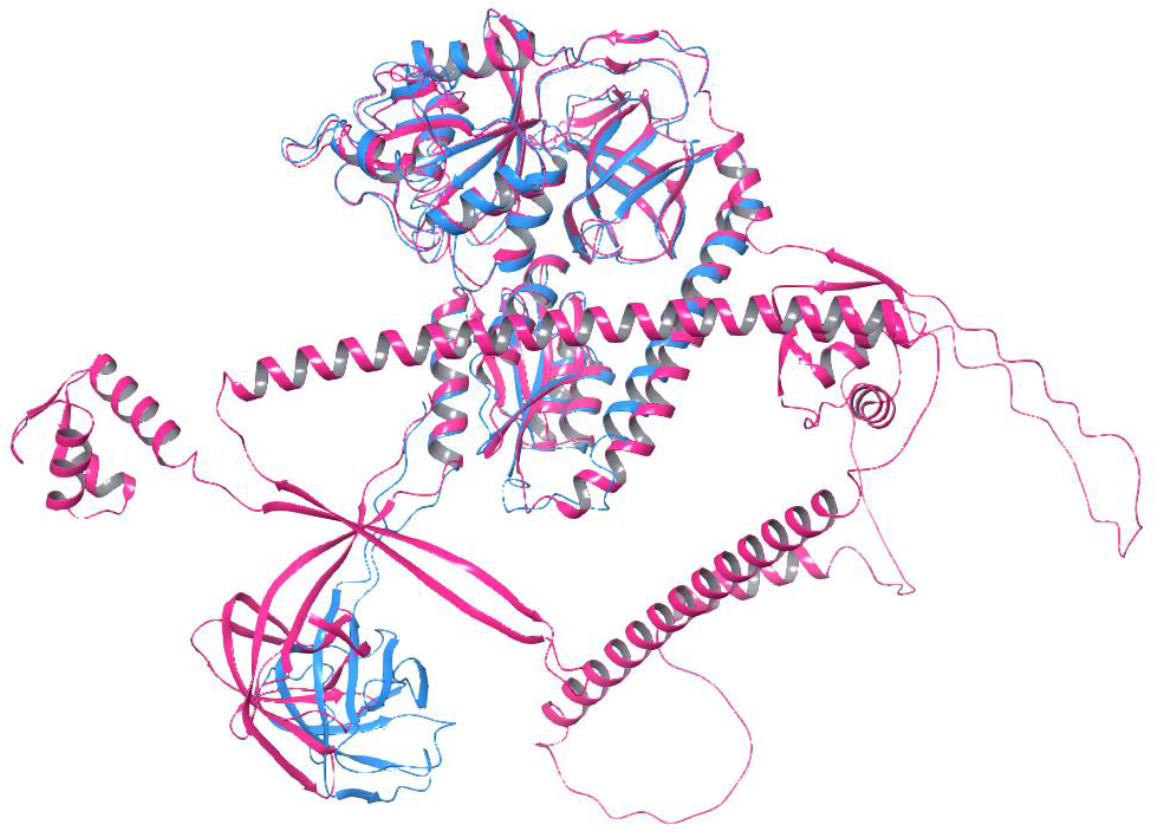
Protein sequence alignment of both IF2 retrieved from UniProt (blue colour) and AlfaFold (magenta colour) at RMSD 2.072

### Reliability of Protein Structure and Prediction of Active Sites

The quality and the reliability of the target protein IF2 in the 3D form is an essential parameter for drug discovery. The Ramachandran plot was obtained to overlay allowed and disallowed zones (Figure 3A), where the newly constructed IF2 protein residues in allowed and favored region thus implying the reliability of the protein structure. The newly constructed IF2 protein was further confirmed for its flexibility and stability. The conformational changes seen in most of the domains. In addition, B-factor of respective domains (I, III and VII) were too high (Figure 3B). This indicates the binding site in these regions will have weak conventional hydrogen bonding contacts upon longevity molecular dynamics that might weaken the binding affinity strength.

**Figure 3:**
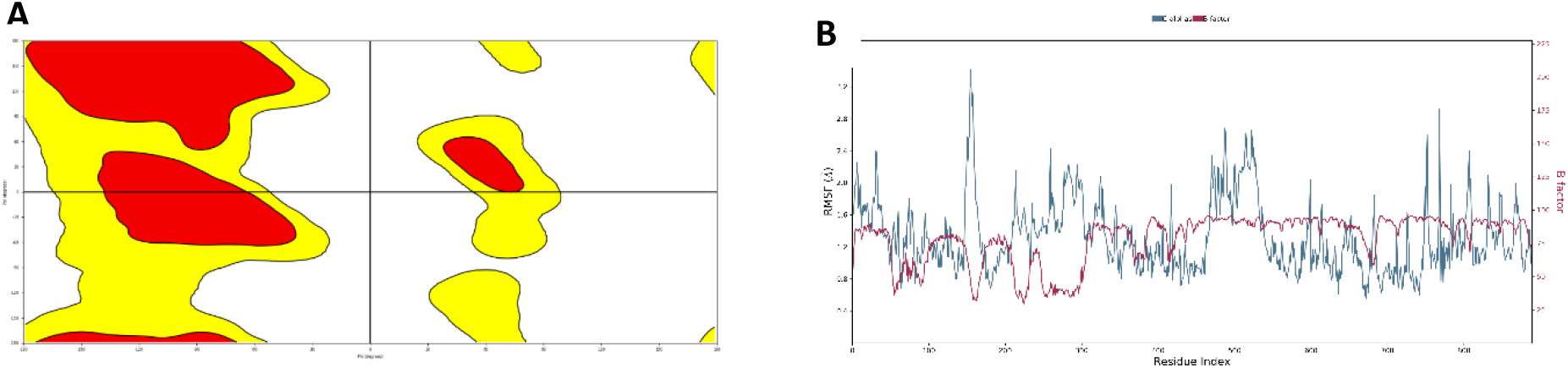
(A) Ramachandran Plot representing the reliability and quality of IF2 protein structure. The yellow is the most allowed region and the reddish is the most favoured region (B) Flexibility and conformational stability of IF2 protein residues

Initially we mapped the different domains regions of IF2 protein based on its residual range (Figure 4). The mapping of various regions ensures us to identify the predicted binding site cavities in designated domains either in shallow or buried regions.

**Figure 4:**
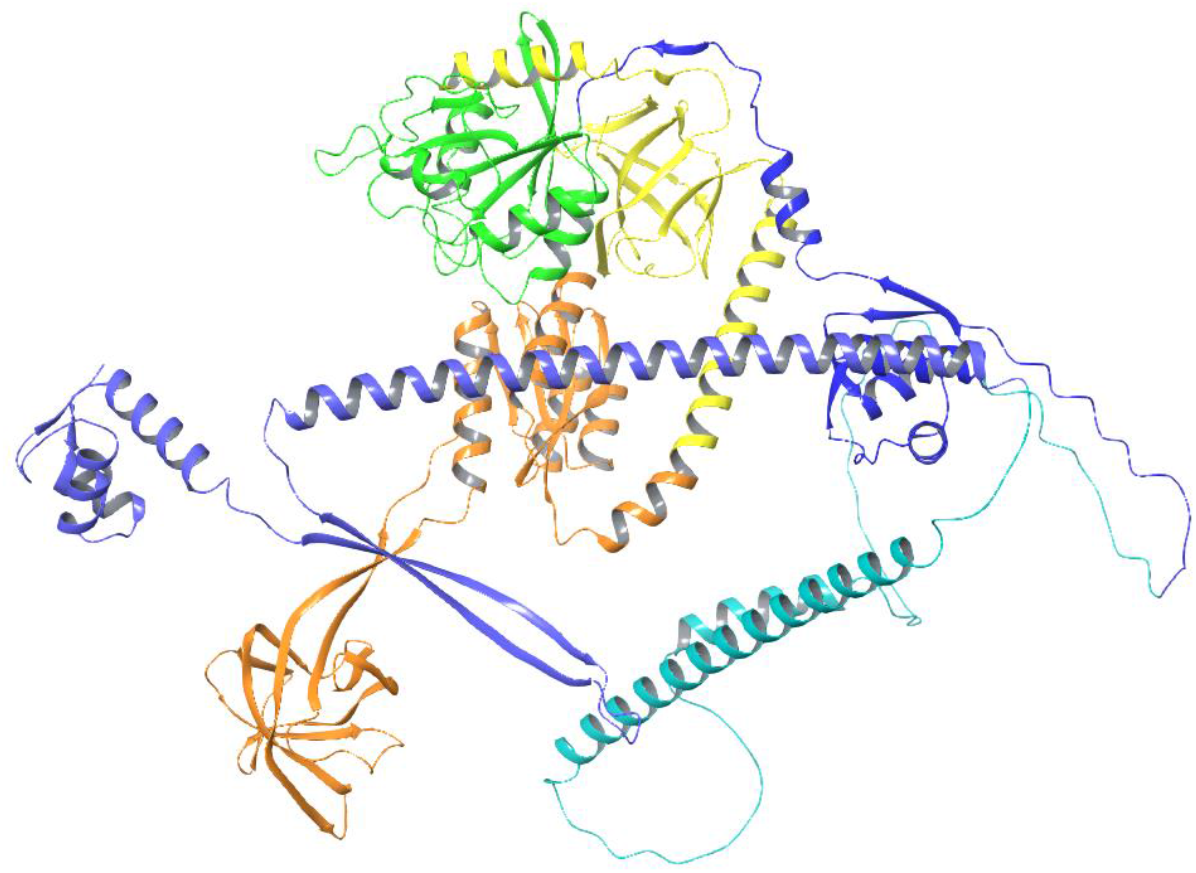
IF2 protein structure representing various domains. The domain colours represent violet (1-158 residues), indigo (159-290 residues), blue (291-392), green (393-540), yellow (541-672) and orange (673-890).

The predicted active binding sites of IF2 using SiteMap module represented in Figure 5A. The representation of the hydrophilic, hydrophobic, HBA, HBD are annotated with the region color as shown in the Figure 5B and 5C. Along with the predicted ones, we precisely pre-defined the domain II with ranging residues (159-290) from the support of the rationale study (Stokes et al., 2014). To consider the promising predicted sites, site score based on DScore (druggability score), surface area and volume needs to be considered for the ligand binding. The site score of 1 or near to 1 or above 1 confirms the high druggability score. The results of all predicted sites are represented in table. From the tabulation, promising site 1 and domain II site were considered as targeted sites for further LTG docking studies. Other sites were not considered because of its narrower cavity and volume size that could led challenges for LTG contacts. Secondly, on the basis of the molar mass of LTG, drug binding at this region will create difficulties in generating surface cavity size and having precise inhibitory affinities. The residues involved with these two sites were extracted with help of Atomic Sequence Language (ASL) within defined radius where LTG could effectively bound (Table 1 and 2).

**Figure 5.**
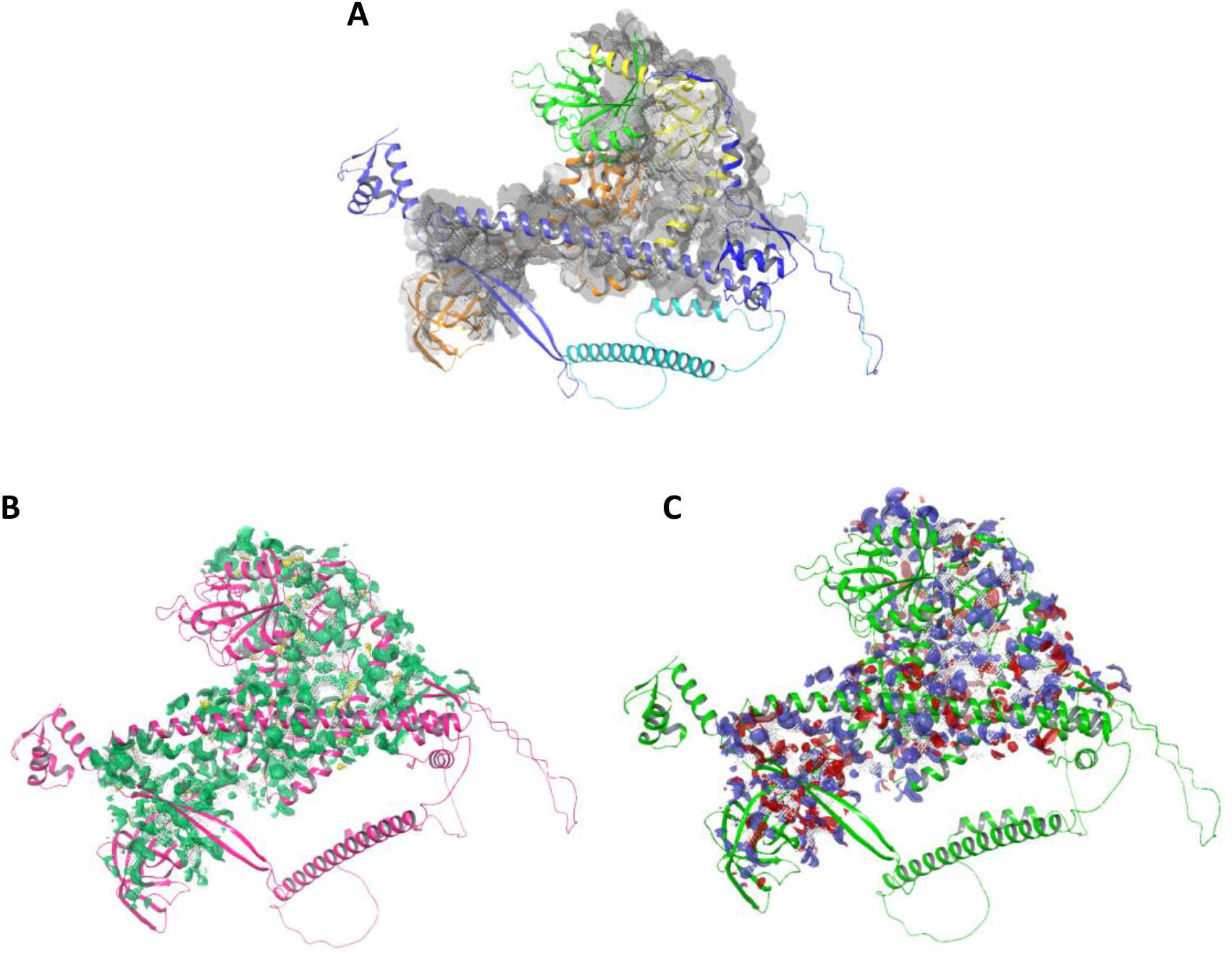
(A) The top-ranked active sites of the protein in greyish regions (B) Magnification of the regions represents hydrophilic regions in green and hydrophobic in yellow (C) Hydrogen-bond donors (blue regions) and hydrogen-bond acceptors (red regions).

**Table 1:** Detailed descriptions of all predicted binding sites of IF2 protein

| Site No. | Binding Site Score | Cavity Size | DScore | Volume |
| --- | --- | --- | --- | --- |
| Site 1 | 0.997 | 105 | 0.934 | 230.153 |
| Site 2 | 0.984 | 126 | 1.011 | 373.870 |
| Site 3 | 0.882 | 77 | 0.922 | 179.046 |
| Site 4 | 0.824 | 56 | 0.823 | 140.973 |
| Site 5 | 0.781 | 47 | 0.774 | 143.374 |
| Domain II | 0.861 | 256 | 1.001 | 74.431 |

**Table 2:** Residue defined from the ASL of top best predicted sites

| Site No. | Residues Involved |
| --- | --- |
| Site 1 | 399, 400, 401, 402, 403, 404, 405, 406, 409, 410, 414, 415, 417, 418, 423, 424, 425, 426, 445, 446, 447, 448, 451, 498, 499, 501, 502, 534, 535, 536, 772 |
| Domain II | 175, 178, 179, 181, 182, 183, 185, 186, 187, 188, 189, 190, 191, 192, 193, 194, 195, 196, 197, 198, 200, 201, 203, 204, 205, 207, 208, 210, 211, 214 |

### Lamotrigine binding to the predicted binding sites of IF2 protein

The docking studies were performed with the best potential predicted site I and the domain II of residues 159-290. For conformational studies, the prepared ligand LTG were simulated using flexible ligand with standard precision at the designated cavities (site I and domain II). Promisingly, we could prove our assertion and the predicted binding site that the binding affinity of LTG exhibited promisingly at site I. The resultant docking score with potential energy of -6.032 kcal/mol. In contrast the docking results also compared with the other sites too to have cross validation of the relative potential energy difference among all predicted sites. From the generated ASL residual sequence at this site Thr425, Gly447 possess strong conventional hydrogen bonding at the triazine moiety of -NH2 and N atom respectively. In addition, the halogen bonding been seen with the -Cl atom of phenyl nucleus projecting towards His 399 and His 402 (Figure 6).

**Figure 6:**
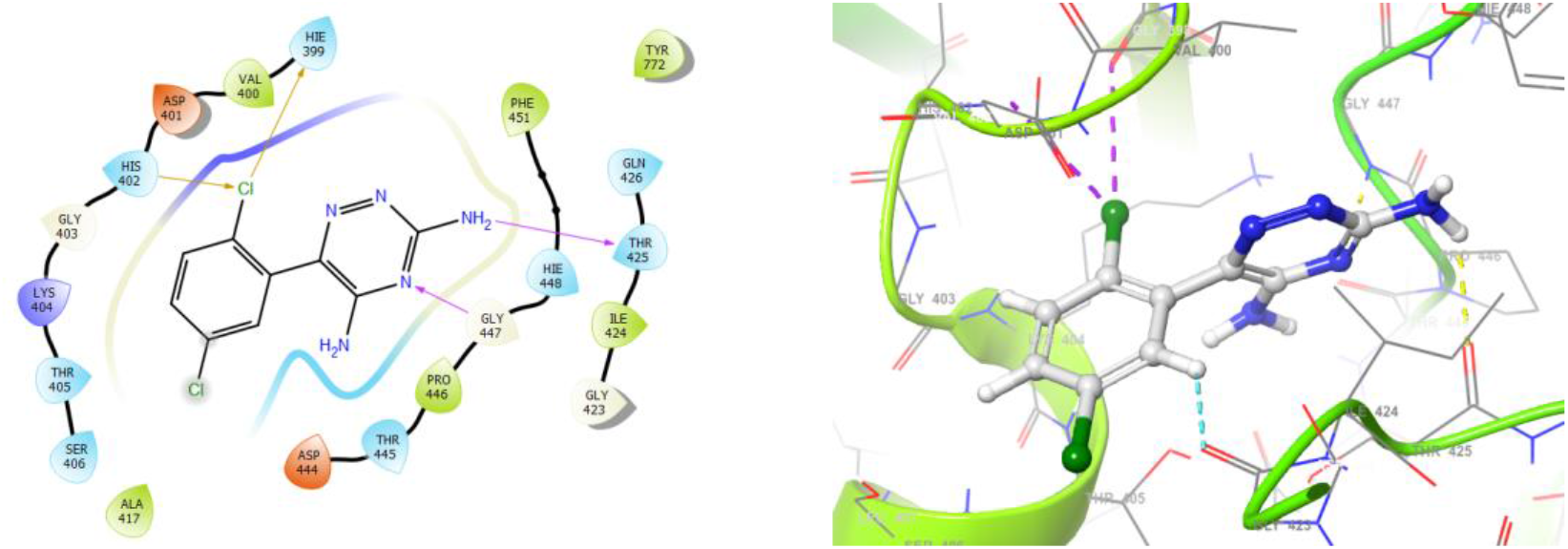
Molecular overlay of LTG binding at predicted site I. Magenta represents hydrogen bonding and orange represents halogen bonding.

The molecular behaviour of LTG at domain II were inclusively studied for conformational analysis as annotated by the Stokes and his co-workers (Stokes et al., 2014). The LTG binding at the ASL sequence range of domain II resulted strong affinity with Glu182 projecting with -NH2 group of triazine nucleus with stronger binding affinity (figure 7). Moreover, the formation of salt bridge with Lys186 projecting towards triazine nucleus might be the synergistic affinity of LTG with IF2 protein. However, the potential force field energy at this domain was -3.472 Kcal/mol due to the binding of LTG at helical region, salt bridge formation and absence of the surrounding residue that could support LTG for binding affinity enhancement at same domain.

**Figure 7:**
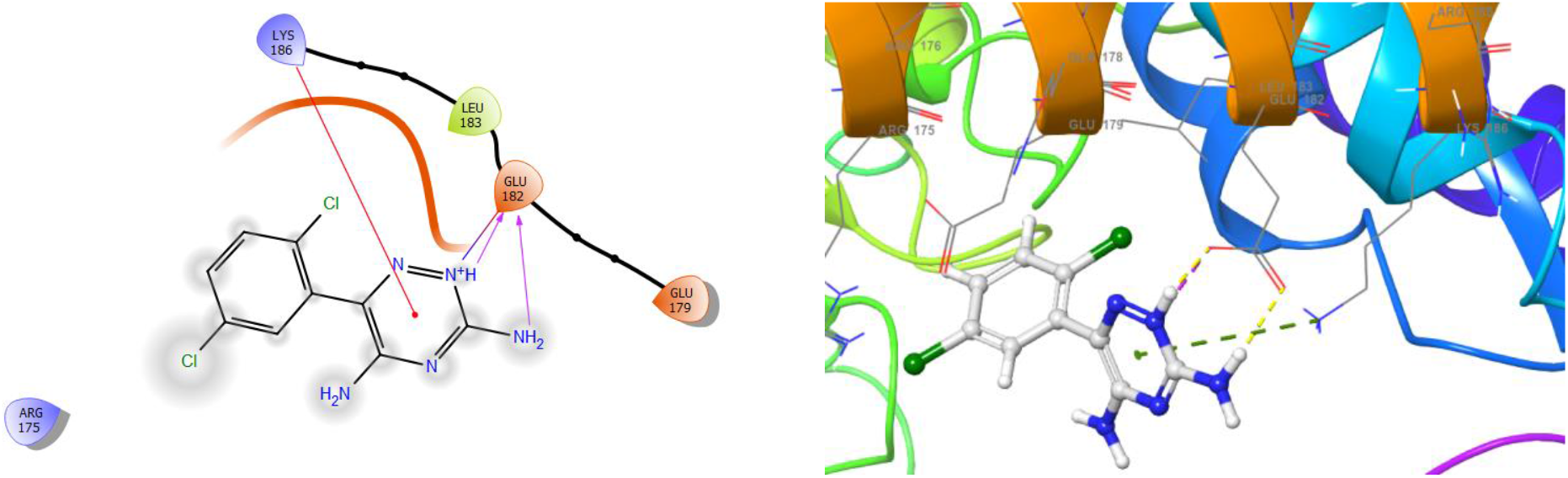
Molecular overlay of LTG binding at domain II. Magenta represents hydrogen bonding and orange represents halogen bonding.

To gain deeper insights and understanding about the molecular behaviour of LTG with the promising predicted site and at domain II, we performed the adaptive dynamics studies at atomic level of bounded LTG crystal with both protein complexes. The RMDS (Figure 8) and RMSF (Figure 9) trajectories suggested us the changes in the cartesian coordinates of the LTG bound with protein on its initial run (up to 20 ns), further could able to stabilize till 50ns. This proves our opinion of the protein flexibility and stability as detailed earlier.

**Figure 8:**
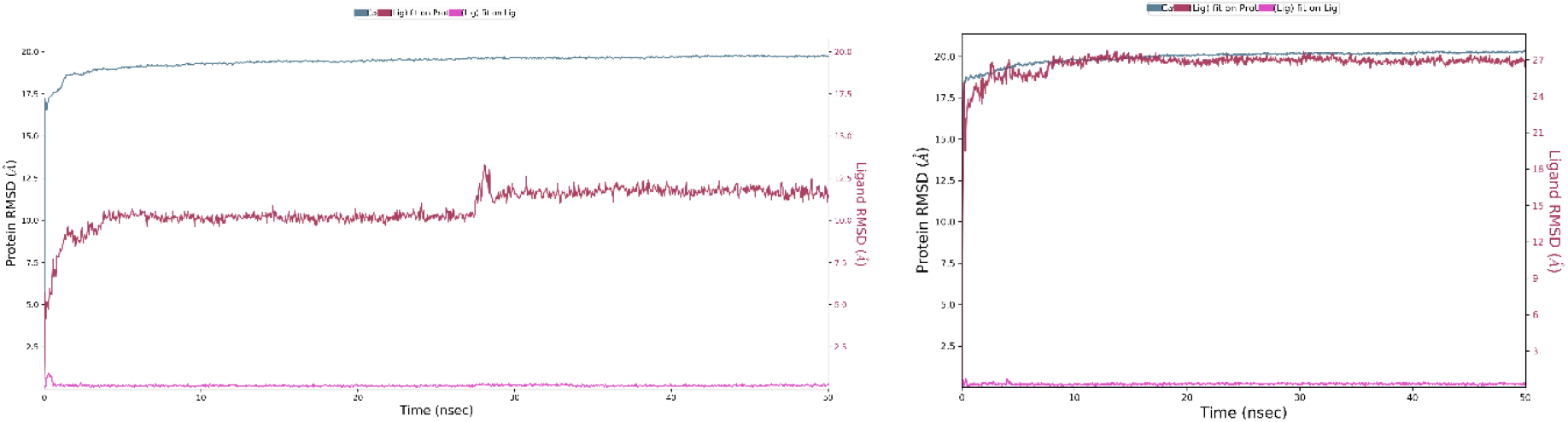
Trajectories representing the LTG bound with (A) site I and (B) domain II of IF2 protein. Green graph represents the protein backbone and red graph is of LTG bound with protein.

**Figure 9:**
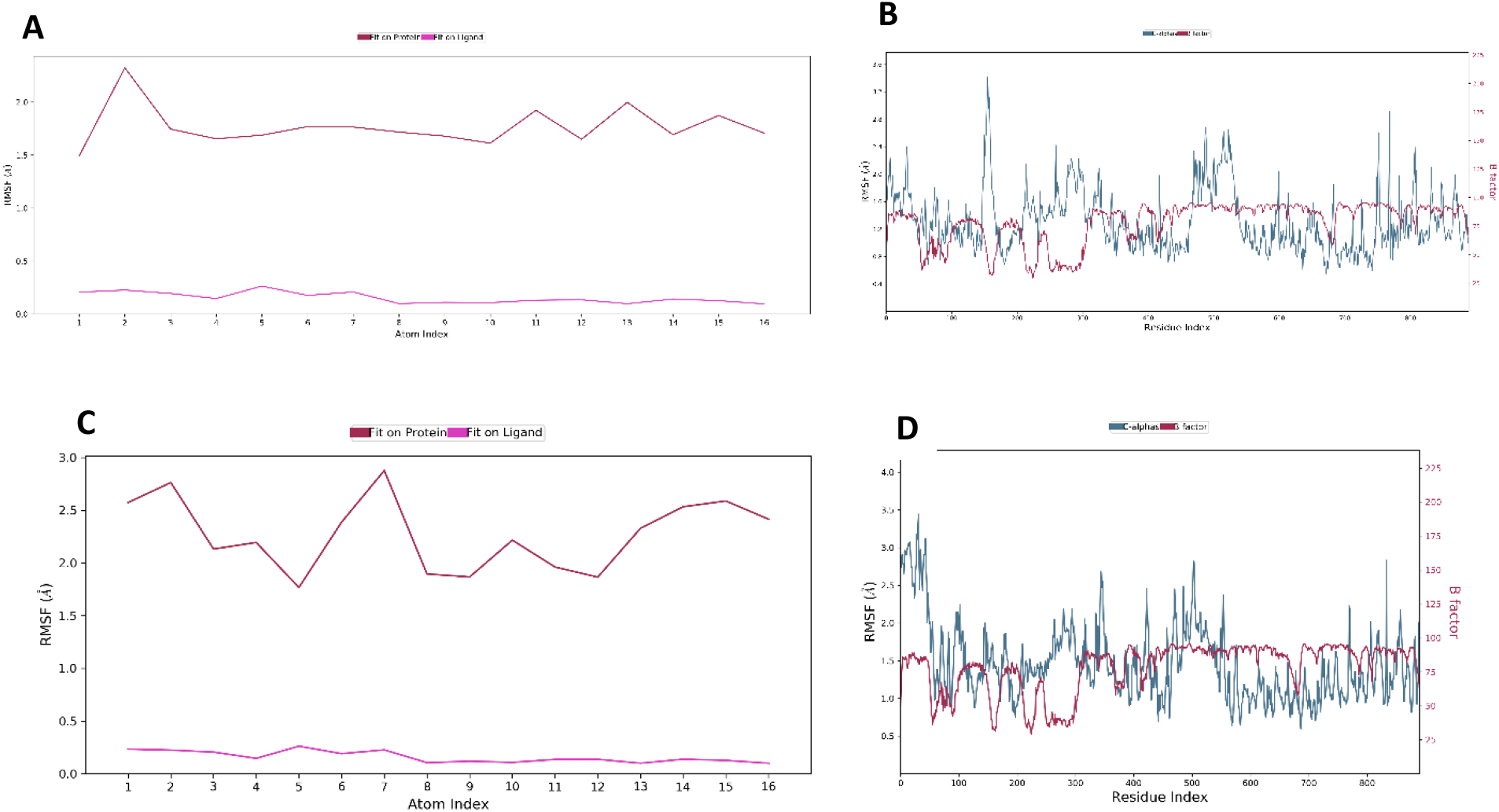
Trajectories representing (A) RMSD of LTG-IF2 and (B) RMSF of LTG-IF2 of site I and (C) RMSD of LTG-IF2 and (D) RMSF of LTG-IF2 of domain II

From the interactions behavioural studied of LTG binding, we found inflation in protein refolding and contacts deviations in contrast with the LTG docked with IF2 protein at 50 ns of simulation studies. As earlier we described due to the high flexibility and spontaneous conformational changes of c-α protein backbone of IF2 protein, the fluctuations of hydrogen bonding contacts could be observed. In detail, the promising active residues for the LTG at respective regions been overlayed with having higher interaction fractions (figure 10). At binding site, I, Thr425 and Tyr772 having 0.9 and 0.5 integration fraction of ligand contacts. In addition, Pro446 and Tyr 772 have higher hydrophobic contacts stability of 0.7 and 1.4 fractions respectively (Figure 10 A and 11A). Similarly, with the domain II site strong hydrogen bonding Arg134, Glu179, Asp751 and Arg755 and hydrophobic contacts with Tyr767 shown the higher interacting fractions (figure 10B and 11B). Here we could see the contradiction in between protein ligand contacts because in the docking studies we found the Glu182 possess higher affinity but upon the simulation runs, we found there been inflation observed in conformational changes of protein backbone that could led failure of protein ligand bonding stability. The reasons could be the presence of domain II (159 – 290 residues) at the helical region and absence of surrounding residues that could strengthen LTG bindings. Secondly the protein flexibility studies (Figure 3B) were too high in this region, where the interactions of LTG with the protein couldn’t be stabilised for longer duration of molecular dynamics simulation studies. Both of these interaction fractions are been calculated in percentage in binding strength after the full simulation runs (figure 12).

**Figure 10:**
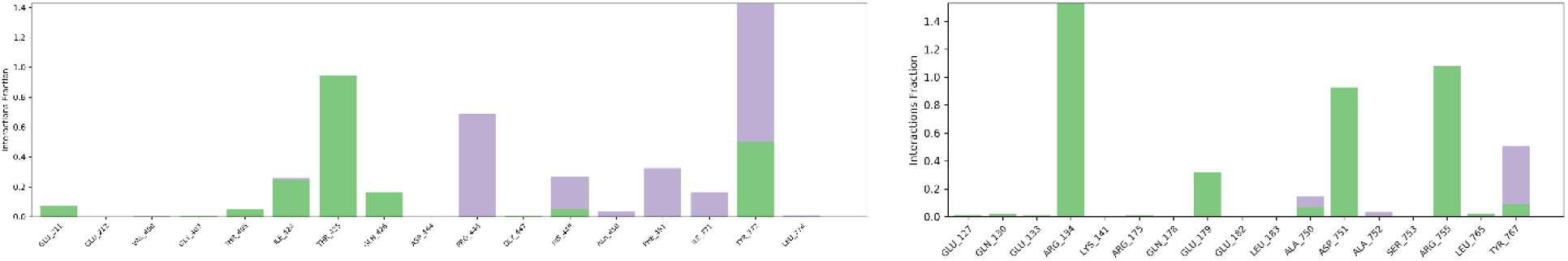
LTG contacts with (A) site I and (B) domain II of IF2 protein represented with its integration fractions. Green represents hydrogen bonding and violet represents hydrophobic contacts.

**Figure 11:**
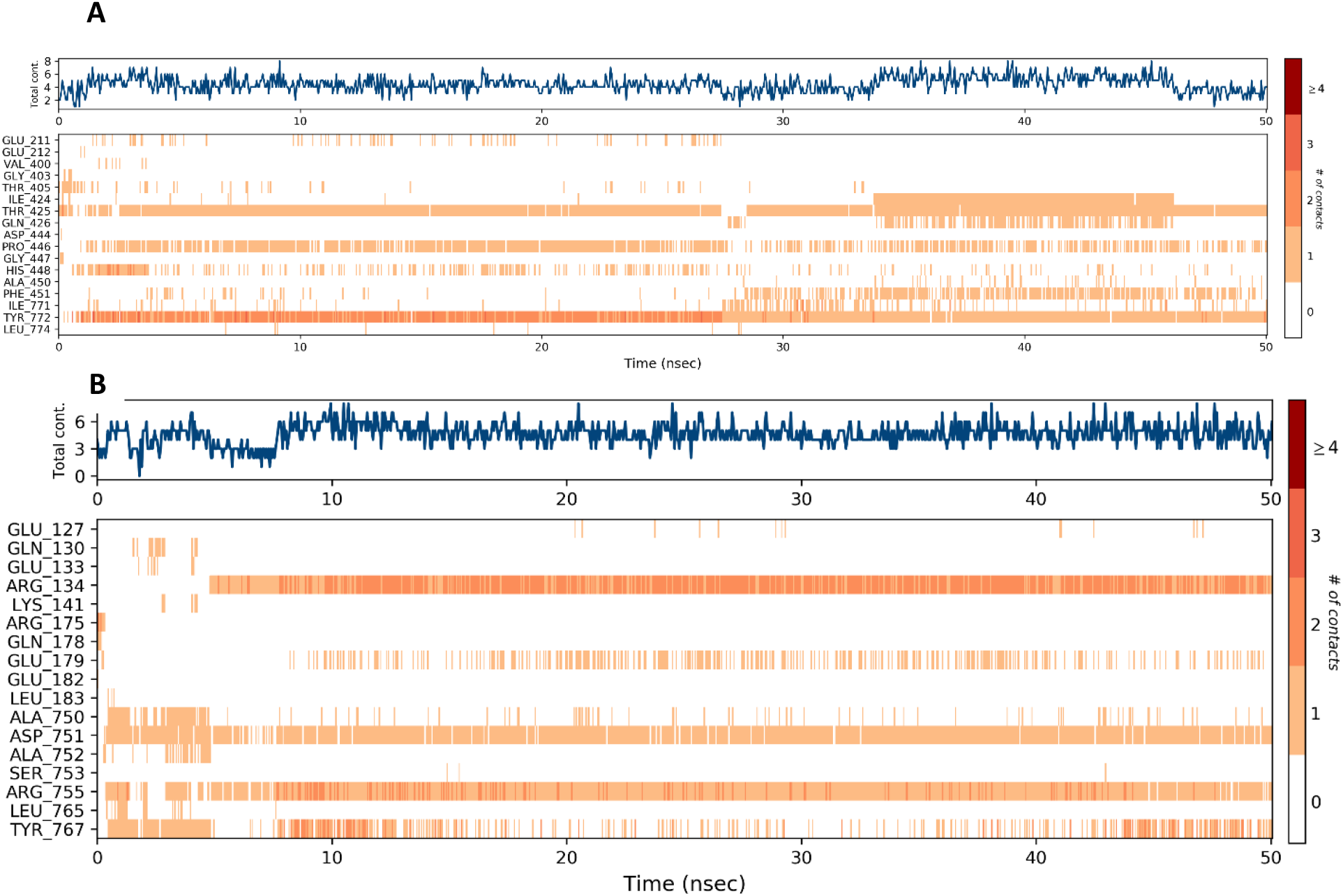
Protein-Ligand interactions timelines representing LTG bounded with (A) site I and (B) domain II of IF2 protein

**Figure 12:**
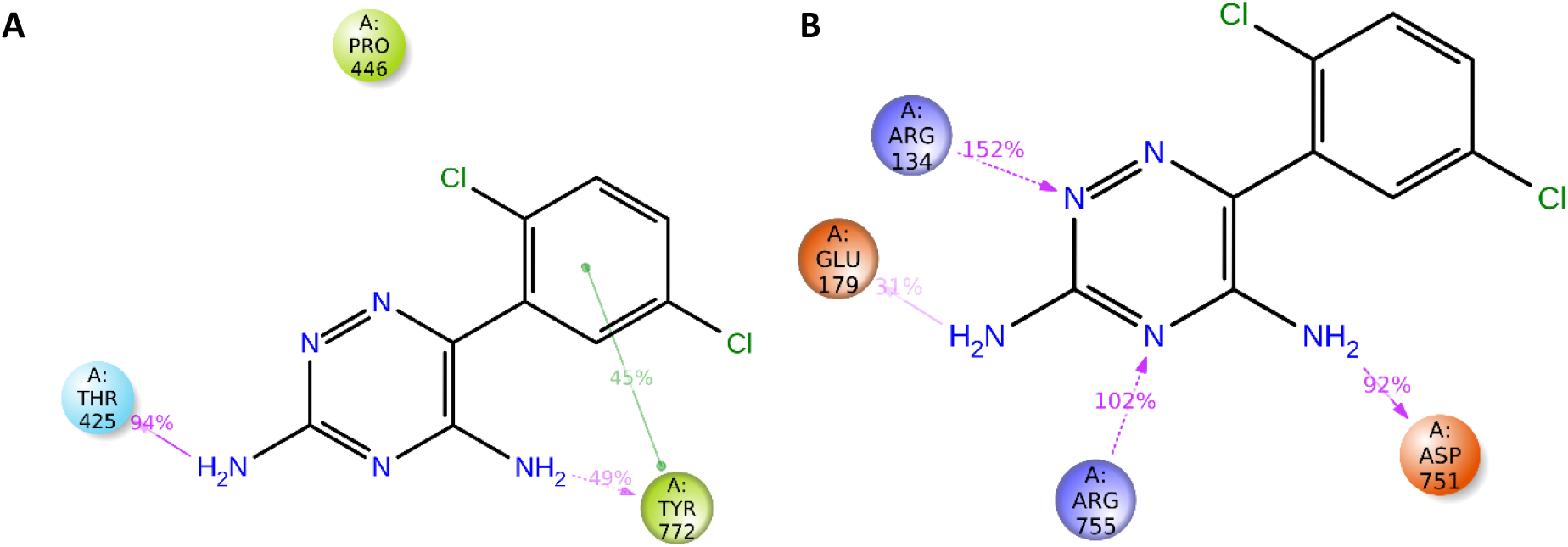
Binding strength of LTG bound with promising residues at (A) site I and (B) domain II of IF2 protein

In our research study, we took an attempt to study the putative residue(s) involved within active binding site cavity of best druggable site predicted (site I) and domain II chain based on the rationale proposed earlier (Stokes et al., 2014). We intensively studied and gained deeper understandings how LTG binds with IF2 protein at atomic and molecular level. With the support of Sprink and his team, we could able to proceed ahead with our structural insights with help of published cryo-EM structure (T. Sprink et al., 2016). The reported cryo-EM structure PDB 3JCN, exhaustively complexed with over 50 proteins comprising IF2, 30s, 50s, mRNA, tRNA, 16s RNA and 23s RNA. Due to partial presence of IF2 protein sequence, we performed synthetic construct with help of AlphaFold database (PDB POA705), which was predicted and structurally aligned with help of powerful AI predictions. The could able to resolve the partial IF2 protein chain and filled missing backbone. We further tried to replace newly constructed IF2 protein in cryo-EM structure for our further study. Further we could able to perform intensive research on the newly designed IF 2 protein and derived the conclusion on its flexibility and conformational stability. Based on the rational study and proven assay studies, LTG able to show bioactivity towards IF2 protein at domain II (Stokes et al., 2014). We further tried to understand the potential residues involved behind the LTG activity with help of computational support. The SiteMap module and CASTp helped us to understand the plausible binding cavity in the whole IF2 protein. In addition, we manually pre-defined with help ASL at domain II falling at range of 159-290 residues.

Gaining deeper understanding and analysis from docking-dynamics results, the proposed assertion of LTG domain II founds to be inaccurate. The reasons could be the presence of intense protein flexibility on the IF2 protein from AlphaFold database, Secondly, binding of LTG sat helical regions led to the weakening of interaction stability. The deviations of contacts from Glu182 falls to Glu179 and addition of other residues projecting towards domain VI (673-890) of IF2 protein. Promisingly, Glu179 could be the crucial residue at domain II for the LTG bioactivity that proven in the assay studies performed by Stokes and his team. The predicted site I (399-772) sequence reveals to be the most promising site, because of the lesser flexibility and the binding of LTG with both alpha and beta sheets. Overall, the interaction seems to be strengthened in contrast with the other predicted sites in same protein.

The totipotency of the LTG at site I results could be considered for further study for multiple proteins of whole IF2 complex and point of randomizations studies for enhancing its synergistic inhibitory activity against multiple bacterial strains and bacterial diseases.

## AUTHOR CONTRIBUTIONS

Both S.A. and J.P. written, performed the computational research and analyzed the data; S.A. conceived the research idea and edited the manuscript.

